# Isolation, Characterization, and Identification of Nitrogen Fixing Bacteria from Rhizosphere of *Sonneratia apetala* Collected from Sundarbans

**DOI:** 10.1101/2024.11.09.622791

**Authors:** Arifa Afrose Rimi, Fatiha Islam Roshnee, Kumar Shuvro, Md. Habibur Rahman Tasin, Rima Akter, Md Emdadul Islam, Kazi M Didarul Islam, S M Mahbubur Rahman, Md Morsaline Billah

## Abstract

Nitrogen is a crucial element for plant growth, driving vibrant greenery, photosynthesis, and overall vitality. This research focused on isolating, characterizing, and identifying nitrogen-fixing bacteria from the rhizosphere of *Sonneratia apetala* in the Sundarbans, Khulna, Bangladesh, with the goal of assessing their potential as biofertilizers. Despite the Sundarbans’ rich microbial diversity, less than 5–10% of species have been identified due to challenges in culturing, which has limited exploration of their applications. In this study, nitrogen-fixing bacteria were isolated using nitrogen-free media, including yeast extract mannitol agar(YEMA) and Burks media, followed by an ammonification test to select ammonia-producing bacteria. This process yielded ten nitrogen-fixing bacterial isolates capable of producing indole-3-acetic acid (IAA). Various biochemical tests, including oxidase, catalase, methyl red, indole, urease, citrate, triple sugar iron, and starch hydrolysis, were conducted. The isolates, designated AK1 to AK10, were identified as *Rossellomorea* sp., *Clostridium* sp., *Achromobacter* sp., *Pseudomonas* sp., *Gluconacetobacter* sp., *Scytonema* sp., *Pseudomonas* sp., *Nesterenkonia* sp., *Gluconacetobacter* sp., and *Bacillus* sp., respectively. Additionally, isolates AK1, AK3, AK4, and AK10 were confirmed through 16S rRNA sequencing. A pot test further revealed that isolate AK-1 significantly stimulated the growth and development of maize seedlings. Future studies are needed to examine the impact of these bacterial isolates on crop yield and seed quality, to better determine their suitability as biofertilizers.

## INTRODUCTION

Bangladesh is grappling with severe overpopulation, leading to an increase in demand for food while cultivable land is continuously diminishing [1]. In response to these challenges, fertilizers are essential for enhancing soil fertility to sustain food production and meet the growing needs of the population. However, chemical fertilizers pose significant environmental and health risks due to the toxic residues they leave behind [2]. Biofertilizers, on the other hand, present a promising alternative, offering a safer and more sustainable approach to boosting soil fertility and crop yield [3–5].

The Sundarbans, the largest mangrove forest in the world, is a unique ecosystem that integrates terrestrial and marine environments and harbors a rich and distinctive array of microorganisms [6]. However, the phylogenetic and functional diversity of microbial communities in mangrove ecosystems remains underexplored compared to other habitats [7]. Among the flora, *S. apetala*, a mangrove species in the family Lythraceae, is prevalent in the Sundarbans and hosts abundant microbial populations in its rhizosphere. These microorganisms are known to play a crucial role in promoting plant growth and contributing to nutrient cycling within the soil [8, 9].

Mangrove ecosystems are shaped by dynamic biological, physical, and chemical processes, influencing nutrient fluxes and supporting the resilience of these habitats [10]. Tidal action, for instance, facilitates the export of mangrove plant detritus, resulting in minimal nitrogen loss from the ecosystem (approximately 3.7 g N per year), equating to around 13% of the average annual net primary production in mangrove forests [11]. Some microbial strains from the Sundarbans rhizosphere have been shown to secrete growth-promoting hormones like auxins, gibberellins, and cytokinins, which are especially beneficial in the high-salinity and anaerobic soil conditions typical of mangrove environments [12].

This study aims to explore the potential of bacterial strains from mangrove ecosystems as biofertilizers to enhance plant growth. While many farmers in Bangladesh rely on chemical fertilizers, such as urea, due to limited awareness of the benefits of nitrogen-fixing bacteria, no nitrogen-fixing bacterial inoculants are available commercially [13]. Developing effective biofertilizers derived from locally abundant nitrogen-fixing bacteria could offer a sustainable solution for improving agricultural productivity.

To address this, we pose the following research question: *Can nitrogen-fixing bacteria isolated from the rhizosphere of S. apetala serve as effective biofertilizers to promote plant growth in Bangladesh’s agricultural systems?* We hypothesize that these nitrogen-fixing bacteria will enhance plant growth by producing growth-promoting compounds like IAA, offering a viable alternative to chemical fertilizers.

The objectives of this study are to isolate nitrogen-fixing bacteria from the rhizosphere soil of *S. apetala*, assess the IAA production potential of each isolate, and characterize and molecularly identify the bacterial species. Ultimately, this research aims to evaluate the impact of these isolates on plant growth, laying the groundwork for their use as biofertilizers.

## MATERIALS AND METHODS

### Soil Collection, Processing, and Bacterial Isolation

Soil samples were collected from the rhizosphere of *S. apetala*in the Sundarbans, using sterile equipment, including polythene bags, spatulas, and gloves. At the collection site, the surface soil was removed to access samples at approximately 6 cm depth. Samples were transferred into sterile 50 mL conical tubes and stored at 4 °C until processing [14].

Soil samples were processed using serial dilution. Initially, 1 g of soil was suspended in 10 mL of distilled water. From this suspension, 1 mL was transferred to a test tube containing 9 mL of sterile water, and this step was repeated across six test tubes. The first tube contained a 1:10 dilution, while subsequent tubes contained further serial dilutions [15]. The bacteria were cultured on YEMA and Burk’s medium to target *Rhizobium* species from the Sundarbans [16]. Ten colonies with distinct morphologies were selected and designated as AK1 to AK10 for further study. Microscopy was used to examine the isolates, which were then incubated for 3–5 days to obtain pure cultures. These cultures were stored with 50% glycerol at -20 °C [17].

### Ammonification Test

The ammonification test, based on the protocol by [18], was used to assess nitrogen-fixing capabilities. Each bacterial isolate was incubated in peptone water for 72 hours, after which Nessler’s reagent was added. A yellow color change indicated the presence of nitrogen-fixing bacteria. Optical density (OD) was measured at 30, 90, and 120 minutes, and a standard IAA production graph was created using OD at 530 nm [19].

### Morphological and Biochemical Characterization

Colony morphology was analyzed on nutrient agar plates after 3–6 days of incubation at 28 °C. Colony characteristics, including color, shape, edge, elevation, and texture, were documented. Gram staining was conducted to classify the isolates. Smears were prepared, heat-fixed, stained with crystal violet for 30 seconds, rinsed, treated with Gram’s iodine for 30 seconds, and washed. A 95% alcohol solution was applied for decolorization, followed by a rinse and counter-staining with safranin for 20–30 seconds. Stained slides were examined under a compound microscope using 10X, 40X, and 100X objectives, with Gram-positive bacteria appearing violet and Gram-negative bacteria pink [20].

For bacterial characterization, various biochemical assays were performed: Catalase, Oxidase, Citrate, Urease, Triple Sugar Iron, Starch Hydrolysis, Indole, Methyl Red, and Nitrate Reduction tests [21].

### Molecular Identification

Bacterial genomic DNA was extracted manually using a lysis buffer with RNAase, proteinase K, chloroform: isoamyl alcohol, isopropanol, and ethanol. DNA samples were stored at -20 °C in 1X TE buffer. DNA quality was verified with 1% agarose gel electrophoresis, and extracted DNA was used in PCR with universal primers targeting the 16S rRNA gene [22]. Sequencing was performed at the National Institute of Biotechnology (NIB), Bangladesh, on a SeqStudioTM Genetic Analyzer, and sequence data were analyzed with BLAST-NCBI, DNA Baser, and MEGA software [23].

### Evaluation of Plant Growth Promotion by Bacterial Inoculation

A nutrient broth culture was prepared for each bacterial isolate in 250 mL Erlenmeyer flasks, incubated for three days in a rotary shaker. Optical density of the inoculum and control was measured using a spectrophotometer. For each isolate, 20 mL of bacterial biomass was applied to a pot with a unique bacterial identification number, and 10 mL of biomass was added to sterilized soil in a poly bag for corn seedling growth. The control group received tap water without microbial inoculation. Plant growth was monitored daily from day 15, and root and shoot lengths were measured to assess the effects of bacterial inoculation on growth [24].

### Statistical Analysis

The data were analysed using analysis of variance with the R using libraries like “agricolae” “dplyr” “ggplot”. The differences among the treatment means were examined using the least significant differences test (LSD) at a 5% probability level (P≤0.05), according Steel and Torrie 1990.

## RESULTS

### Sample Collection, Preparation, Bacterial Isolation, and Microscopic Observation

To study colony morphology, a widely-used method for identifying bacterial characteristics on agar, ten isolates with distinct morphologies were identified, as detailed in supplementary table 1 and illustrated in supplementary figure 1 [25]. These isolates successfully grew on selective media, confirming their nitrogen-fixing ability, as illustrated in Figure 1. Colony-forming units (CFU) per mL were calculated using the equation C*FU* = (number of colonies × dilution factor) / culture volume, yielding a colony count of 4.1 × 10 □ CFU/mL. Most isolates showed a red color on Gram staining, indicating a Gram-negative classification, with AK1 and AK10 being the exceptions, showing Gram-positive results [26].

**Table 1.** Biochemical test results of bacterial isolates. Here, the (+) sign indicates a positive and the (-) sign indicates a negative result.

| Isolates | Catalase | Oxidase | Glucose | Lactose | Sucrose | H<sub>2</sub>S | Urease | Methyl red | Indole | Starch hydrolysi | Nitrate reduction | Citrate | Predicted Species |
| --- | --- | --- | --- | --- | --- | --- | --- | --- | --- | --- | --- | --- | --- |
| AK1 | + | + | + | + | + | + | - | - | - | + | + | + | *Rossellomorea sp.* |
| AK2 | - | - | - | + | - | - | - | - | - | + | - | - | *Clostridium sp.* |
| AK3 | + | + | - | - | - | - | + | - | - | - | + | - | *Achromobacter sp.* |
| AK4 | + | + | - | - | - | - | + | - | - | + | - | + | *Pseudomonas sp.* |
| AK5 | + | - | + | + | + | - | - | - | - | - | - | - | *Gluconacetobacter sp.* |
| AK6 | - | - | - | - | - | - | - | - | - | - | - | - | *Scytonema sp.* |
| AK7 | + | - | + | + | + | - | - | - | - | + | + | - | *Pseudomonas sp* |
| AK8 | + | - | - | - | - | - | + | - | - | + | - | - | *Nesterenkonia sp.* |
| AK9 | + | - | - | + | + | - | - | - | - | + | - | - | *Gluconacetobacter sp.* |
| AK10 | + | - | + | + | + | - | - | - | - | - | + | + | *Bacillus Pumilus* |

**Figure 1.**
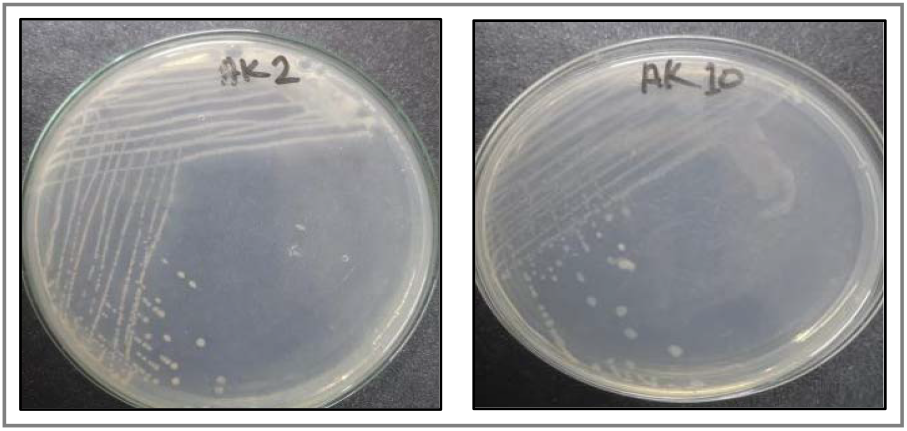
Bacterial culture in Burk’s media. Where AK1 and AK10 indicates the bacterial culture number.

### Ammonification Test

All isolates produced ammonia, evident by a color change from yellow to shades of orange or red depending on the amount produced. Notably, isolates AK1 and AK10 displayed the highest ammonia production, as indicated by optical density measurements at 530 nm. Using the equation Y = 0.0027x - 0.0043, the calculated concentrations for AK1 and AK10 were 66.4 mg/mL and 52.3 mg/mL, and the rest of the isolates concentration are given in supplementary Table 2 and figure 2 [27].

**Table 2.** The Genetic Information of highest result showing bacterial isolates (depending on plant growth and IAA production) from NCBI, consisting of query length, similarity and reference number.

| Internal code | Name of bacteria | Accession Number | BLAST match sequence |  |  |
| --- | --- | --- | --- | --- | --- |
|  |  |  | Reference number | Query length | Similarity (%) |
| AK1 | *Rossellomorea aquimaris* | OQ407551.1 | *Rossellomorea aquimaris* KR709236.1 | 1299bp | 99.54% |
| AK3 | *Achromobacter xylosoxidans* | OQ407542.1 | *Achromobacter xylosoxidans* LC125142.1 | 319bp | 99.37% |
| AK4 | *Stenotrophomonas geniculata* | OQ407541.1 | *Stenotrophomonas geniculata* MG571680.1 | 1275bp | 99.22% |
| AK10 | *Bacillus Pumilus* | OQ407541.1 | *Bacillus Pumilus* MN642632.1 | 1290bp | 99.77% |

**Figure 2.**
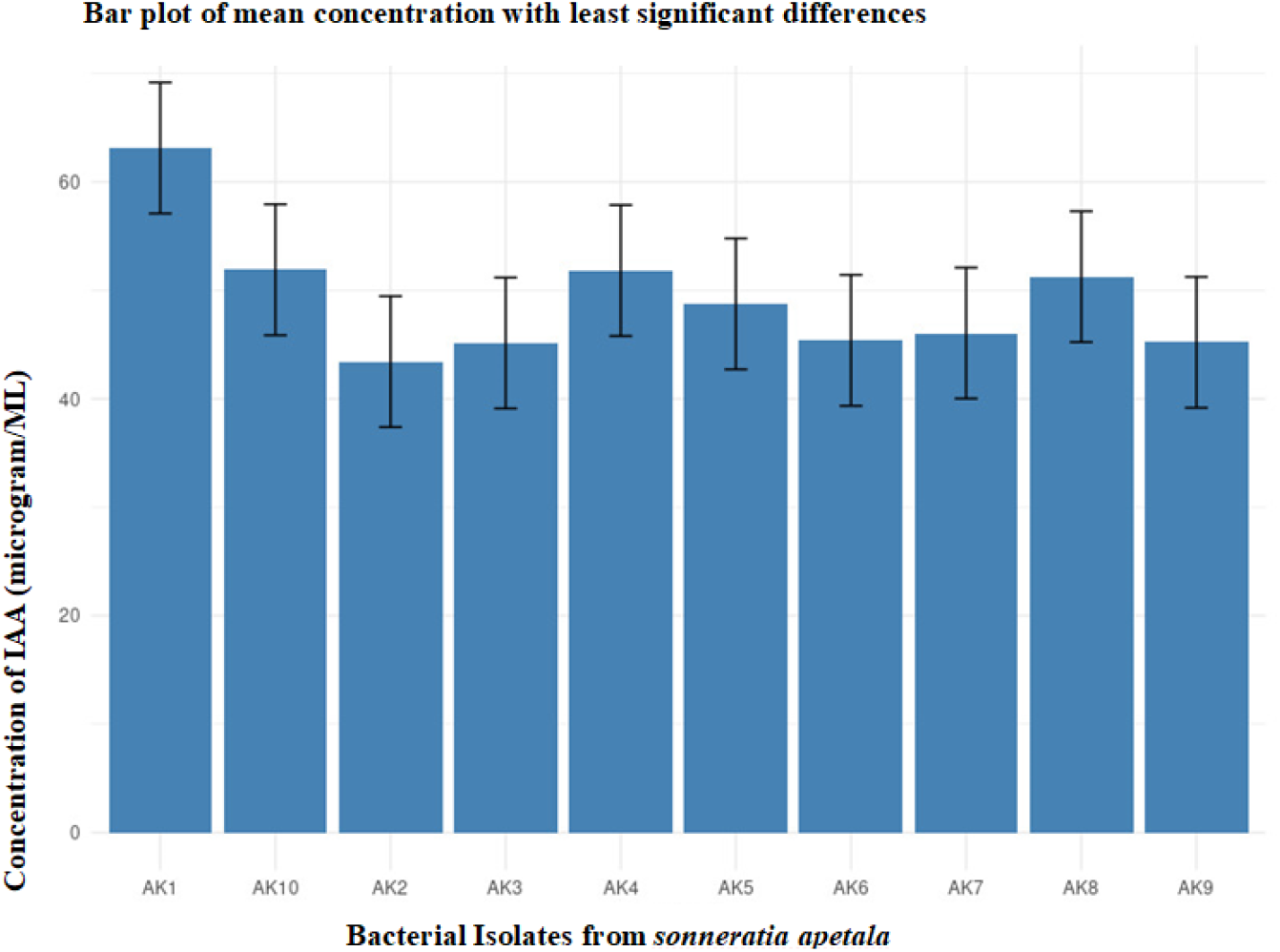
IAA concentration graph. Where n=3, the error bars represent the minimal significant difference between treatments at P < 0.05.

### Biochemical Tests

The biochemical and physicochemical test results are presented in Table 1. Among the isolates, AK1, AK3, AK4, and AK10 exhibited positive results for citrate utilization, oxidation, catalase, and nitrate reduction tests, with AK10 showing unique results in certain tests. All isolates except AK10 tested negative for the methyl red test, while all isolates were negative for the indole test. In the urease test, isolates AK3 and AK4 showed positive results but were negative in the TSI test. In contrast, isolates AK1 and AK10 yielded positive results for TSI but negative for urease, with these reactions indicated by color changes [28]. Based on these biochemical findings, the isolates AK1–AK10 were preliminarily identified as *Rossellomorea sp*., *Clostridium sp*., *Achromobacter sp*., *Pseudomonas sp*., *Gluconacetobacter sp*., *Scytonema sp*., *Pseudomonas sp*., *Nesterenkonia sp*., *Gluconacetobacter sp*., and *Bacillus sp*., respectively [29].

### Molecular Identification

DNA extraction and PCR were conducted on all ten samples, and the presence of DNA was confirmed through gel electrophoresis. To verify the identities of isolates AK1, AK3, AK4, and AK10, sequencing was performed, taking into account the IAA production and plant growth rates. The 16S rRNA sequencing results identified AK1 as *Rossellomorea aquimaris*, AK3 as *Achromobacter xylosoxidans*, AK4 as *Pseudomonas geniculata*, and AK10 as *Bacillus pumilus*. Genetic data for these isolates are available from the NCBI database, as listed in Table 2 and the constructed phylogenetic tree is shown in figure 3 [28, 30].

**Figure 3.**
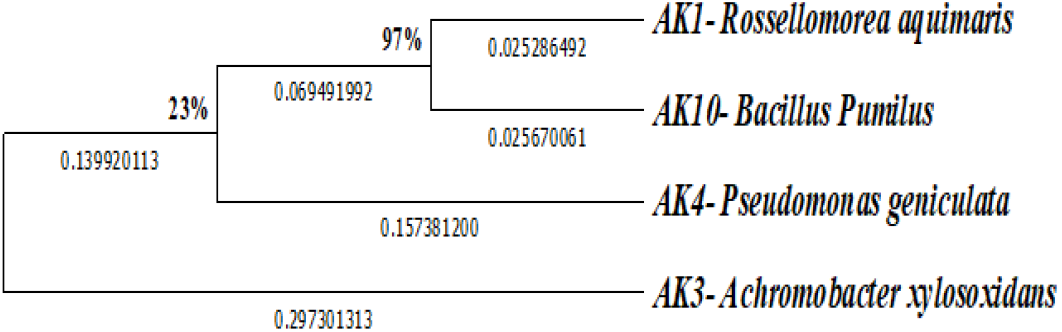
Taxonomic Evolutionary relationships, and phylogenetic tree. The ancestral lineage was determined through the application of the Neighbor-Joining method. (NeiM.1987) The best-fit tree is displayed beneath the branches, and the calculation of evolutionary distances employed the Maximum Composite Likelihood method [60]. The provided information pertains to a phylogenetic analysis involving four nucleotide sequences. The evolutionary distances, measured in the number of base substitutions per site, are depicted in a tree diagram. The fraction of places where at least one unambiguous base appears in at least one sequence for each descendant clade is shown next to internal nodes. The analysis considered codon positions, including 1st, 2nd, 3rd, and noncoding positions, with the removal of all ambiguous positions in each sequence pair using pairwise deletion. The final dataset comprised 1325 positions, and the evolutionary analyses were carried out using MEGA11.

### Plant Growth Analysis

To examine the impact of bacterial isolates on plant growth, a controlled pot experiment was conducted at Khulna University. Each pot received 20 mL of microbial biomass from individual isolates, and an additional 10 mL was applied to sterilized soil in polyethylene containers. Plants treated with bacterial inoculants demonstrated enhanced root and shoot growth compared to the untreated control group [31]. Among the inoculated plants, those treated with isolate AK10 showed the most substantial growth, with AK10 producing 52.3 μg/mL of both ammonia and IAA, respectively. Isolate AK1 also contributed positively to plant growth, with an IAA concentration of 66.4 μg/mL, exhibiting superior overall plant growth metrics compared to the other isolates. Detailed results are provided in supplementary table 3 and supplementary figure 2 also illustrated in Figure 4.

**Figure 4.**
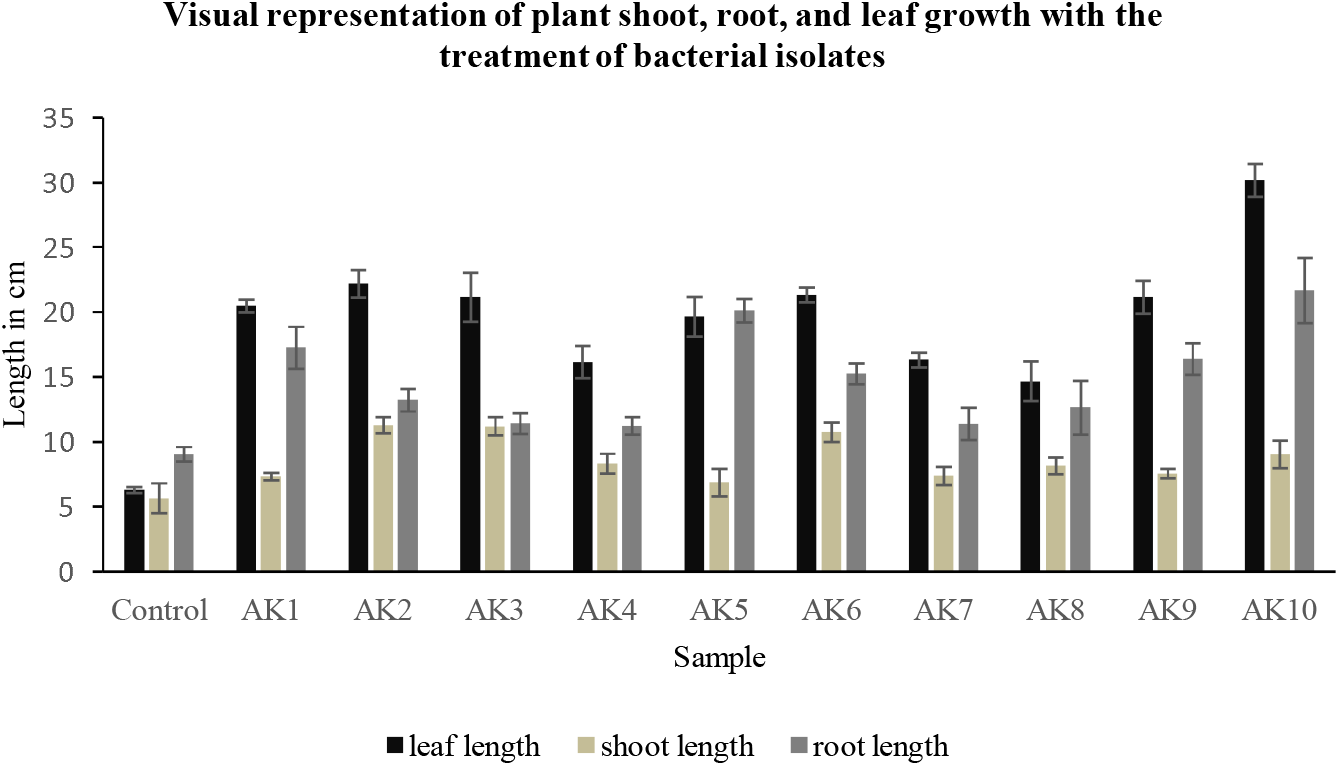
Impact of bacterial inoculation on the lengths of shoots, roots, and stems of maize plants cultivated in growth pouches under axenic circumstances. Where n=3 and the error bars indicate the minimum significant difference between treatments at P ≤ 0.05.

## DISCUSSION

Research on the spatial distribution of microbial diversity in mangrove ecosystems is still relatively limited [32]. This study investigated the nitrogen-fixing ability of bacteria from the rhizosphere of *S. apetala*, a key mangrove species collected from the Sundarbans Forest in Khulna, Bangladesh. The comprehensive characterization of bacterial isolates was conducted to assess their potential as biofertilizers, offering a sustainable alternative to synthetic fertilizers with minimal environmental impact [33]. The Sundarbans, known for its rich microbial diversity, hosts a variety of nitrogen-fixing bacteria that could enhance plant growth [34].

In our study, soil samples from the rhizosphere of *S. apetala* were processed using serial dilution, and bacteria were cultured on selected media. Ten isolates (AK1–AK10) were identified based on colony morphology and various biochemical tests, including ammonification and IAA production. All isolates were confirmed to produce ammonia, with isolates AK1 and AK10 exhibiting the highest levels. For precise identification, four isolates (AK1, AK3, AK4, and AK10) underwent sequencing using 16S rRNA. The pot experiments demonstrated that the bacteria, particularly AK1 and AK10, significantly enhanced maize growth, indicating their potential as biofertilizers to replace synthetic nitrogen sources.

IAA plays a crucial role in plant growth by promoting root initiation, cell division, and overall growth, which enhances the root surface area and improves access to soil nutrients [35-36]. Our findings confirmed that the nitrogen-fixing bacteria isolated from the rhizosphere of *S. apetala* exhibited significant plant growth-promoting properties, including nitrogen fixation and IAA generation. Among the isolates, four species demonstrated notable growth enhancement effects on maize plants.

The isolation methods employed in this study were effective, echoing findings from previous research that successfully cultured nitrogen-fixing bacteria like *Rhizobium* from the Sundarbans region [16]. Our study isolated ten bacterial strains from genera such as *Rossellomorea, Clostridium, Achromobacter, Pseudomonas, Gluconacetobacter, Scytonema, Nesterenkonia*, and *Bacillus*. The identification of AK1 as *R. aquimaris*, AK3 as *A. xylosoxidans*, AK4 as *P. geniculata*, and AK10 as *B. pumilus* was confirmed through 16S rRNA sequencing.

The sequencing analysis revealed that AK1 shared 99.7% similarity with *R. aquimaris*, a species recognized for its plant growth-promoting capabilities, including phosphate solubilization and IAA production. Strains of *R. aquimaris* have been previously isolated from the rhizosphere of tomato plants and are known to enhance nutrient availability, improve plant growth, and support sustainable agricultural practices [37]. Isolate AK3, with 98.7% similarity to *A. xylosoxidans*, is known for fixing atmospheric nitrogen in various crops, which improves productivity [38-39]. Previous studies have linked the activities of *A. xylosoxidans*—including phosphate solubilization and IAA synthesis—to enhanced growth in crops like wheat and rice [40-41]. Isolate AK4, identified as *P. geniculata*, belongs to the family Pseudomonadaceae and is recognized for its role in forming beneficial interactions with plant roots. *Pseudomonas* spp. is integral to plant development, nutrient accumulation, and stress responses in the rhizosphere [42]. The strain *P. geniculata* produces and bioaccumulates soluble sugars, proline, and antioxidant enzymes, aiding maize plants in salt stress tolerance and promoting growth [46]. Among the isolates, AK10, identified as *Bacillus pumilus*, is particularly noteworthy. This strain promotes plant growth through the production of IAA, 1-aminocyclopropane-1-carboxylic acid deaminase, cytokinins, gibberellins, and contributes to atmospheric nitrogen fixation and phosphorus solubilization [47].

In vitro screening revealed that the isolated bacteria produced IAA in concentrations ranging from 45.29 to 66.41 μg/mL, indicating significant diversity in IAA production among *S. apetala* rhizosphere isolates. Notably, isolates AK1 and AK10, producing 66.41 μg/mL and 52.33 μg/mL of IAA, respectively, demonstrated exceptional potential as growth regulators [48]. This finding aligns with earlier studies highlighting the plant growth-promoting effects of rhizobacteria in mangrove habitats [48]. The observed variation in IAA production can be attributed to differences in biosynthetic pathways, gene locations, and enzymatic activities [49]. The IAA concentrations in our study were higher than those previously reported, suggesting that the soils studied contain bacteria with highly sought-after properties for plant growth enhancement [50-52].

The pot experiment showed that bacterial inoculation with AK1 and AK10 significantly improved maize plant growth after a 15-day growth period, as evidenced by increased shoot, root, and leaf lengths compared to the control group. This growth promotion can be attributed to the synergistic effects of nitrogen fixation, IAA production, and phosphate solubilization, which are crucial for enhancing nutrient availability in nutrient-deficient soils like those found in the Sundarbans mangrove ecosystem.

While this study presents valuable insights, it acknowledges certain limitations. The varying responses of different bacterial isolates to plant growth can be attributed to their unique traits, a finding supported by consistent auxin production results from previous studies [34, 53]. Future research should expand beyond the focus on IAA to explore other phytohormones and their combined effects on plant growth [54]. The controlled conditions of our experiments, although essential for understanding plant-bacterial interactions, may not fully replicate real agricultural environments, highlighting the need for further field trials [55]. The study did not cover the entire microbial diversity of the rhizosphere, which limits the understanding of nutrient cycling; however, the use of 16S rRNA sequencing ensured precise identification of bacterial strains [56]. Additionally, while the correlation between root length and nitrogen uptake is promising, validation across different crops and conditions is necessary [57].

To enhance the robustness of our findings, we employed three replicates for each biochemical assay, from sample collection to execution, ensuring accuracy. Selective media were used to limit the growth of non-target microorganisms. Reagents and equipment were sourced from reputable suppliers to ensure precision in our methods. Although biochemical tests are useful for identifying and characterizing bacterial strains, they have limitations in distinguishing closely related strains. Therefore, we recommend full-length 16S rRNA sequencing for accurate species identification [58-59]. The strong root and shoot growth activity observed with *R. aquimaris* suggests its potential for biofertilizer applications.

To advance research on nitrogen-fixing bacteria from the rhizosphere of *S. apetala* in the Sundarbans, several priorities should be established. Testing these strains on diverse crops and conducting field trials in various environments will be essential for assessing their broader effectiveness. Developing bacterial consortia that combine beneficial traits such as nitrogen fixation, phosphate solubilization, and hormone synthesis (IAA, gibberellins, cytokinins) may lead to improved plant growth outcomes. Further research should also explore the combined effects of phytohormones and utilize metagenomics to understand the full microbial community present in the rhizosphere. Investigating strains like *R. aquimaris* for phytoremediation could further support environmental restoration efforts. Optimizing biofertilizer formulations and conducting long-term impact studies will provide insights into sustained benefits, while genetic research may enhance strain capabilities. Collaborating with local farmers will ensure the practical relevance of our findings and promote sustainable agricultural practices.

## CONCLUSION

This study has successfully identified a diverse array of bacterial communities within the rhizosphere of *S. apetala*, highlighting the presence of specific strains capable of nitrogen fixation. The isolation of these nitrogen-fixing bacteria was effectively accomplished using nitrogen-free selective media, followed by validation of their nitrogen-fixing abilities through ammonification tests. Additionally, the production of IAA was assessed, further enhancing the potential of these isolates as biofertilizers. Comprehensive characterization of the bacterial isolates included gram staining, colony morphology assessment, and a series of biochemical tests, all conducted in triplicate to ensure reliability and precision.

Subsequently, the selected bacterial isolates were applied to the soil during the seed-sowing phase of a pot experiment. After a 15-day growth period, significant differences in growth parameters, including plant height and dry weight, were observed when compared to the control group. These findings underscore the potential of these bacterial strains to promote plant growth, which is crucial for sustainable agricultural practices.

However, further research is imperative to fully understand and optimize the biofertilizer potential of these isolates. Future studies should focus on assessing their impact on crop yield and seed quality, particularly in the agricultural context of Bangladesh, where such enhancements could significantly benefit the economy. Given the importance of agriculture in emerging economies, this research contributes valuable insights toward developing sustainable agricultural practices that harness microbial diversity for improved agricultural productivity.

## ACKNOWLEDGMENTS

We would like to express our sincere gratitude to the National Institute of Biotechnology (NIB), Bangladesh, for their invaluable support and assistance with the sequencing of the bacterial isolates in this study. Their expertise and resources significantly contributed to the successful completion of our research. We also thank all colleagues and staff of Biochemistry and Molecular Biology Laboratory, Biotechnology and Genetic Engineering Discipline, Khulna University who facilitated our work and provided insightful discussions throughout the project.

## COMPLIANCE WITH ETHICAL STANDARDS

This article does not contain any studies with human participants performed by any of the authors.

