## Supplementary file for "Isolation, Characterization, and Identification of Nitrogen Fixing Bacteria from Rhizosphere of *Sonneratia apetala* Collected from Sundarbans"

TABLES

**Supplementary Table 1.** The colony morphology of bacterial isolates

| Sl no | Name | Media | Growth Time (Day) | Morphology | | | | Colony |
|---|---|---|---|---|---|---|---|---|
| | | | | Color | Form | Elevation | Margin | |
| 1 | Ak1 | YEMA | 3 | Orange | Circular | Flat | Undulate | Medium, Smooth |
| 2 | Ak2 | YEMA | 3 | White | Circular | Convex | Entire | Medium, Smooth |
| 3 | AK3 | YEMA | 2 | Creamy white | Irregular | Flat | Entire | Medium, Smooth |
| 4 | AK4 | YEMA | 2 | White | Circular | Flat | Entire | Medium, Smooth |
| 5 | AK5 | YEMA | 2 | Orange | Circular | Flat | Entire | Medium, creamy |
| 6 | AK6 | YEMA | 2 | White | Punctiform | Flat | Erese | Medium, Smooth |
| 7 | AK7 | YEMA | 2 | Orange | Circular | Flat | Entire | Medium, Smooth |
| 8 | AK8 | YEMA | 2 | White | Punctiform | Flat | Undulate | Small, Rough |
| 9 | AK9 | YEMA | 2 | Pale Yellow | Punctiform | Flat | Erese | Medium, Rough |
| 10 | AK10 | YEMA | 3 | white | circular | convex | entire | creamy |

**Supplementary Table 2.** Indole-3 Acetic Acid concentration

| **Name** | **Count** | **Average Concertation of IAA(μg/ml)** |
|---|---|---|
| AK1 | 3 | 63.13 |
| AK2 | 3 | 43.43 |
| AK3 | 3 | 45.14 |
| AK4 | 3 | 51.82 |
| AK5 | 3 | 48.75 |
| AK6 | 3 | 45.39 |
| AK7 | 3 | 46.06 |
| AK8 | 3 | 51.26 |
| AK9 | 3 | 45.20 |
| AK10 | 3 | 51.89 |

**Supplementary Table 3.** Average root, shoot and leaf lengths of maize plant. There is one control and three replications for each bacterial isolate treated plant

| **Groups** | **Count** | **Leaf length average (cm)** | **Shoot length Average (cm)** | **Root length Average (cm)** |
|---|---|---|---|---|
| Control | 3 | 6.3 ± 0.25 | 5.66 ± 1.15 | 9.06 ± 0.55 |
| AK1 | 3 | 20.5 ± 0.5 | 7.33 ± 0.28 | 17.26 ± 1.61 |
| AK2 | 3 | 22.2 ± 1.05 | 11.3 ± 0.60 | 13.23 ± 0.87 |
| AK3 | 3 | 21.16 ± 1.89 | 11.2 ± 0.7 | 11.43 ± 0.81 |
| AK4 | 3 | 16.16 ± 1.25 | 8.33 ± 0.76 | 11.23 ± 0.68 |
| AK5 | 3 | 19.66 ± 1.52 | 6.86 ± 1.05 | 20.13 ± 0.9 |
| AK6 | 3 | 21.33 ± 0.57 | 10.76 ± 0.75 | 15.26 ± 0.8 |
| AK7 | 3 | 16.33 ± 0.57 | 7.4 ± 0.7 | 11.4 ± 1.22 |
| AK8 | 3 | 14.66 ± 1.52 | 8.16 ± 0.66 | 12.63 ± 2.07 |
| AK9 | 3 | 21.16 ± 1.25 | 7.56 ± 0.37 | 16.4 ± 1.21 |
| AK10 | 3 | 30.16 ± 1.25 | 9.03 ± 1.05 | 21.66 ± 2.51 |

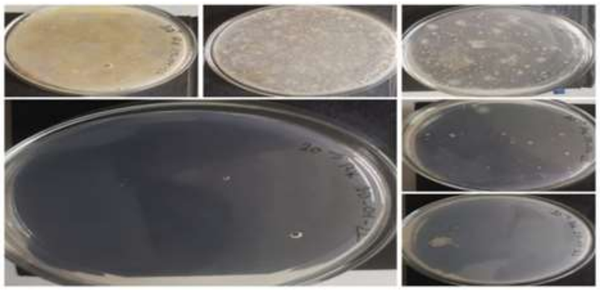

**Supplementary** Figure 1: Bacterial isolation by dilution culture method

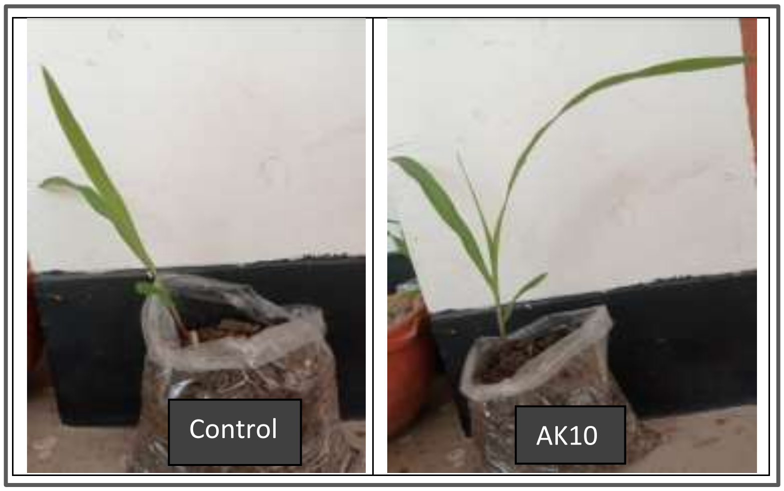


**Supplementary** Figure 2: Pot test-15 days picture of control and AK10
